# Aperiodic exponent of brain field potentials is dependent on the frequency range it is estimated

**DOI:** 10.1101/2024.12.17.628966

**Authors:** Gonzalo Boncompte, Vicente Medel, Martin Irani, Jean Phillip Lachaux, Tomas Ossandon

**Affiliations:** División de Anestesiología, Escuela de Medicina, Pontificia Universidad Católica de Chile; Latin American Brain Health Institute, Universidad Adolfo Ibañez, Santiago de Chile, Chile; Neurodynamics of Cognition Lab, Departamento de Psiquiatría, Pontificia Universidad Católica de Chile; Université Claude Bernard Lyon 1, CNRS, INSERM, Centre de Recherche en Neurosciences de Lyon CRNL U1028 UMR5292, Bron, France

**Author notes:** Correspondance should be addressed to and/or.

**Keywords:** aperiodic exponent, aperiodic slope, 1/f slope, intracortical recordings, Specparam, FOOOF, IRASA

## Abstract

The aperiodic component of brain field potentials, like EEG, LFP and intracortical recordings, has shown to be a valuable tool in basic neuroscience and in clinical applications. Aperiodic activity is modeled as a power law of the power spectral density, with the aperiodic exponent as the key parameter. Part of the interest in this parameter lies in its proposed role as a marker of the balance between excitatory and inhibitory cortical activity. In theory, a perfect power law would imply that the same behaviour exists across all frequencies, however recent evidence has suggested that low and high frequency ranges could present different aperiodic exponents. To elucidate this, we systematically evaluated the relation between frequency range and aperiodic parameters using human resting-state intracortical recordings from 62 patients. We employed two distinct estimation methods, Specparam and IRASA. We found that aperiodic parameters were indeed dependent on frequency range. Specifically, we found that low frequency ranges displayed, on average, lower aperiodic exponents (flatter power spectral density) than high frequency ranges. This behaviour was consistent for Specparam and IRASA estimations in all frequency ranges compatible with EEG. Given that there is currently no consensus for a single frequency range to be used in either clinical or basic neuroscience, our results show that care should be taken when comparing aperiodic exponents derived from different frequency ranges. We believe our results also encourage further research into the possible roles that aperiodic exponents estimated from different frequency ranges could have in reflecting distinct aspects of cortical systems.

## Introduction

Neural field potential signals can be decomposed into oscillatory and aperiodic activity (Donoghue et al., 2020). Oscillations are the result of coordinated increases and decreases in underlying neural activity that occur with a (relatively) regular period (Buzsáki et al., 2012; Steriade, 2005). In contrast, background aperiodic activity does not occur in a particular frequency but throughout the whole spectrum, or at least a significant portion of it. The power spectral density (PSD) of electroencephalographic (EEG), intracortical and local field potential (LFP) recordings consistently shows an inverse relation between power and frequency whereas, in general, low frequency components have greater power than high frequency components. For at least 25 years now (Pereda et al., 1998), this relation between power and frequency has been modeled as a power law, following equation (1):

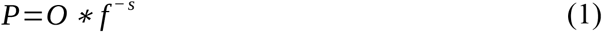

where *P* is spectral power, *f* is each frequency component and *O* and *S* are free parameters, aperiodic offset and slope respectively. The slope parameter, also called aperiodic exponent or 1/f slope, has gained considerable interest recently, partly because it has been linked to a specific neurobiological property, the balance between excitatory and inhibitory activity in cortical (Gao et al., 2017) and subcortical areas (Wiest et al., 2023). This is consistent with empirical results showing higher slopes, meaning a greater difference in power between low and high frequency components, are found during anesthesia, sleep or generally inhibitory cortical states (Medel et al., 2023; Zhang et al., 2023). On the other hand, lower slope values, where high and low frequency activity have more similar magnitudes, have been reported in epilepsy (Coa et al., 2022), attentional deficit and hyperactivity disorder (Ostlund et al., 2021) and Parkinson’s disease (Mostile et al., 2019; Wiest et al., 2023). This evidence posits aperiodic properties as valuable tools in both clinical and basic neurobiological settings.

The recent increase in studies analyzing aperiodic slope of brain field potentials has showcased an important variability in the values of aperiodic slope, possibly related to how these aperiodic parameters are estimated. Ideal power law behaviour would be fully independent on the frequency range used, e.i aperiodic parameters estimated between 0.1 Hz and 10 Hz or between 30 Hz and 100 Hz should be equivalent (perfectly scale-free). However, empirical power laws are not ideal and at most only follow equation (1) in a particular frequency range (Newman, 2005; Stumpf & Porter, 2012). This has been recognized even by early works that described power-law behaviour on human EEG data (Pereda et al., 1998). In part because of this, aperiodic parameters are estimated using only a specific frequency range within PSDs. Importantly, although the choice of frequency range is deemed very important for proper estimation (Gerster et al., 2022; Kramer & Chu, 2023), a general consensus for the criteria for this decision is currently lacking in the literature; frequency range is decided by simple visual inspection of PSDs, to try to avoid known sources of artifacts or other factors. A wide variety of frequency ranges have been employed to estimate aperiodic parameters including 30 – 45 Hz (Lendner et al., 2020), 1 – 40 Hz (Lanzone et al., 2022) and 4 – 50 Hz (Robertson et al., 2019) (see Table 1 in Kramer & Chu, 2023 for further examples). It has even been proposed that PSDs could be better described by two distinct power-law’s, one on low frequency and one in high frequency ranges (Ferree & Hwa, 2003; He et al., 2010). It has also been suggested that works that estimate aperiodic slope in higher frequency ranges tend to produce higher estimates for aperiodic slope (Kramer & Chu, 2023; Zhang et al., 2023). If proven true, this would raise an important question with regards to the comparability of aperiodic parameters estimated in different frequency ranges. Relatedly, it remains unknown if there are particular frequency ranges which are more stable and thus suitable for aperiodic exponent estimation.

To address these questions, here we analyzed 62 human intracortical recordings obtained during resting state. We systematically studied the relation between frequency and aperiodic exponent using the two complementary analytical methods to estimate aperiodic activity in brain field potentials (Donoghue et al., 2024), namely Specparam, formerly known as FOOOF (Donoghue et al., 2020) and Irregular Resampling Auto-Spectral Analysis (IRASA) (Wen & Liu, 2016). We found that there is indeed a strong positive dependency between frequency and aperiodic exponent. In addition, we found no frequency range in which aperiodic exponent was fully stable. These results are discussed in the context of recent literature.

## Methods

### Study population and Intracortical recordings

Intracranial recordings were obtained from 62 epileptic patients with intractable epilepsy who underwent stereotactic intracerebral EEG (SEEG) recordings before surgery at the Neurology Department of the Grenoble University Hospital (Grenoble, France). The implantation decision was only based on clinical criteria. All electrode data presenting pathological waveforms were discarded from the present study. This procedure was achieved in collaboration with the medical staff. It was based on the visual inspection of the recordings and by systematically excluding data from any electrode sites that were found a posteriori to be located within the seizure onset zone. All participants provided written informed consent, and the experimental procedures were approved by the local Ethical Committee (“ISD et SEEG” project, CPP Sud-Est V no 09-CHU-12).

### Signal preprocessing

Artifact-containing segments, or periods of ictal activity from each patient’s signals were rejected via clinical criteria and visual inspection complemented with an automatic detection algorithm, following a previously validated approach (Kucyi et al., 2020). Signals were re-referenced to the average of all electrodes. Then, from each patient’s data we selected 5 minute clean segments for further processing and analysis. Signals were acquired at either 1024 Hz or 512 Hz. We homogenized all signals to have a common sampling frequency of 512 Hz.

### Power spectral density estimations

Power spectral density (PSD) estimations were conducted in each clean 5 minute window as follows. The signal was segmented into non-overlapping 2 s epochs, after which multitaper spectral estimation was conducted using 7 DPSS windows (Python, MNE) using frequency limits of 0.5 to 255.5 Hz. The PSD estimated for each electrode and subject was the median value across epochs.

### Frequency ranges and spectral widths

To evaluate the possible dependence of aperiodic slope to the frequency range used to estimate it we employed a series of frequency ranges that varied in their Center Frequency (CF) and their spectral width. CF refers to the middle point, in a logarithmic scale, of the frequency range. Spectral width refers to the magnitude of the span of frequencies around the CF and its measured in octaves. Low and high frequency limits for each frequency ranges are calculated according to:

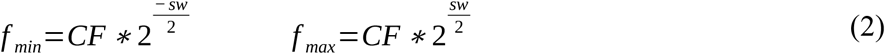

where f_min_ and f_max_ are the minimal and maximal frequencies of a frequency range, CF is the center frequency and *sw* corresponds to the spectral width. This results in that, for a spectral width of 2 octaves, a given frequency range of center frequency = CF_i_ is composed by frequencies ranging from [CF_i_ / 2] to [2 * CF_i_]. We initially choose a base of 55 distinct Cfs that spanned from 3 to 200 Hz for every spectral width. For some center frequencies, this resulted in frequency ranges that went above the Nyquist frequency of our data (256 Hz) or went above those limited by IRASA resampling method limitations. These frequency ranges were omitted.

To evaluate the consistency across subjects of the relation between aperiodic paramenters and CF, we conducted Spearman correlations between groups of 15 aperiodic exponents and 15 contiguous CFs, which were centered around what we called “mid frequency”. In this way we could observe if the correlation between aperiodic exponent and CF was different in different frequencies.

### Specparam (FOOOF)

Specparam algorithm was applied to median PSDs for each electrode and subject. For each PSD we estimated aperiodic slope (exponent), offset and *r*^2^ as a goodness of fit measure. For spectral width of 2 octaves this resulted in the inclusion of 49 frequency ranges. The lower of these went from 1.5 to 6 Hz (CF = 3 Hz) and the higher one went from 62.7 to 250.8 (CF = 125.4 Hz). We did not employ the “knee” option available in Specparam, meaning that just a classical power law was used in the aperiodic model. Default parameters of the FOOOFGroup class were employed. Models that could not be fitted with sufficient accuracy (r^2^ < 0.98) were rejected from further analyses.

### IRASA

IRASA also estimates the aperiodic components of a signal but uses a different strategy than specparam. It relies on the fact that periodic and narrow-band oscillations change their frequency when the original signal is resampled (Wen & Liu, 2016). The IRASA algorithm then calculates PSDs of several resampled versions of the original signal and calculates the median across them. Oscillatory (periodic) activity will be lost in the calculation of the median while components that are aperiodic, i.e. not present only in a narrow frequency band, will remain in the final median. Then the best linear curve was estimated for each median aperiodic-specific PSDs from which aperiodic slope, offset and *r*^2^ values were extracted We employed the Python library YASA, with resampling factors from 1.1 to 1.9 with increments of 0.05 and a time window (2 s).

### iDFT simulations

To evaluate the base effects that both algorithms have (Specparam and IRASA) in a fully controlled setting, we simulated signals using inverse Discrete Fourier Transform (iDFT) model (Medel et al., 2023). These simple models are directly constructed in the frequency domain from the PSD and only then are translated into the time domain using the inverse Fourier transform. In this way we were able to fully control for the spectral properties of the tested signals. Once in the time domain we evaluated the estimated aperiodic parameters using Specparam and IRASA using the same parameters as the ones used in empirical data.

We constructed 6 series of simulations using either one or two power laws in different frequency ranges. All series had a power law with an aperiodic exponent of 1 in the low frequency range (0.5 to 10 Hz) and a second power law with an aperiodic exponent that was different for each series of simulations (0.5, 1.0, 1.5, 2.0, 2.5 or 3.0). This second power law with variable aperiodic exponent was present from 10 to 240 Hz (Figure 3). The data presented for each serie is the average of 100 equivalent simulations.

## Results

To better understand the relation between aperiodic parameters and the frequency range used to estimate them, we calculated aperiodic parameters in 62 open eyes resting state intracortical recordings using a variety of frequency ranges. We systematized our exploration by using center frequencies (CF) ranging from 3 to 125 Hz. Each frequency range had a span of 2 octaves, resulting in frequency limits from half of the CF to twice the CF (e.g. for CF of 30 Hz, frequency limits were 15 to 60 Hz, see Methods).

### Aperiodic parameters versus CF using Specparam

First we estimated the aperiodic slope of each patient’s data using the Specparam algorithm. We found that aperiodic exponent strongly depended on the frequency range used (Figure 1A). Throughout all CF values lower than ∼30 Hz, aperiodic slope increases as CF values increased. Our results show a similar positive dependency between aperiodic offset and CF (Figure 1B). To estimate the degree of consistency of the correlation between aperiodic parameters and frequency ranges used, we calculated a series of Spearman correlations in regularly spaced subsets of 15 CF ranges (Figure 1C, D; see Methods). To illustrate this, ten groups of CFs with different mid frequencies are depicted in each column of Figure 1C and Figure 1D for slope and offset respectively. For example in Figure 1C, the first column (mid frequency of 5.2 Hz) reports the correlation between 15 aperiodic slopes estimated with 15 different CFs (ranging from 3 Hz to 8.9 Hz). Statistical evaluation of these correlations between slope and offset showcased that practically all subjects displayed significant positive correlations at mid frequencies lower than aproximatedly 30 Hz (Figure 1E, F). In higher frequencies a divergent behaviour occurs: subjects show either positive or negative correlations, but very few (∼10%) presented non significant correlations between aperiodic exponent or offset and CF (Figure 1E, F, gray lines).

**Figure 1.**
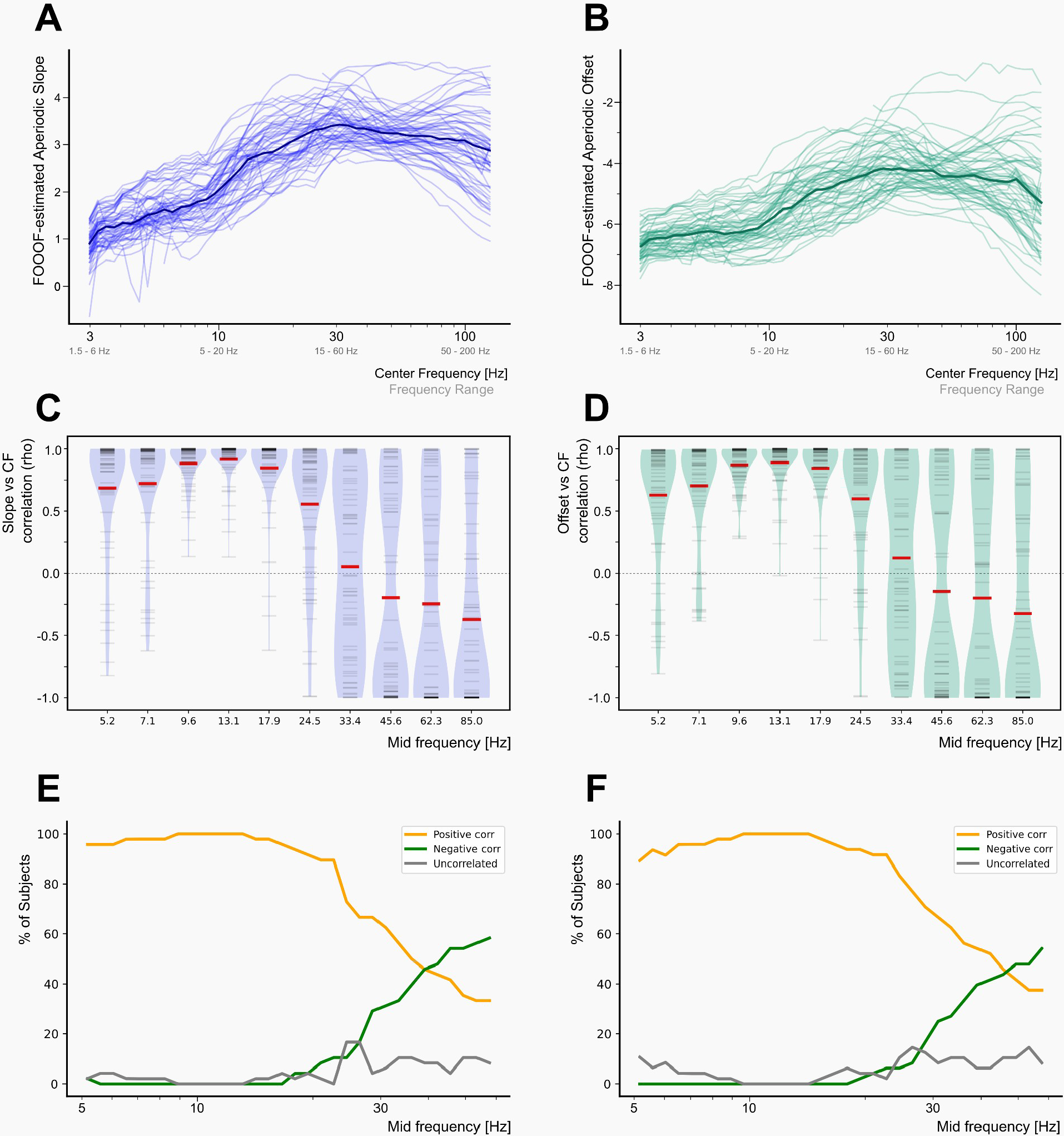
Aperiodic parameters depend on the frequency range in which they are estimated when using Specparam. **(A-B)** Line plot showing, for each subject (soft lines), the aperiodic slope (A) and (B) offset estimated within each frequency range. Dark lines show the median slope and offset across subjects. **(C-D)** Plots illustrating the Spearman correlation coefficients (rho) between aperiodic slope (C) and offset (D) when assessed in different ranges of CFs. Mid frequency refers to the middle frequency of 15 consecutive frequency ranges. Each subject’s rho values, for each mid frequency is represented by a horizontal gray line in each column. Each colum’s violin plot displays the probability density function of rho values across subjects. Median rho values for each mid frequency are represented by a thick red horizontal line in each column. **(E-F)** Line plots showing the percentage of subjects, for each mid frequency, that displayed a significant positive correlation, significant negative correlation or a non significant correlation between aperiodic parameter (slope or offset) and CF. Significance of Spearman correlations was corrected for multiple comparisons.

### Aperiodic parameters versus CF using IRASA

To further evaluate this relation, we conducted an analogous analysis of the dependence of aperiodic slope and offset to CF using the IRASA method. IRASA-estimated aperiodic slope and offset showed a positive dependence towards CFs in the whole range of frequencies tested (Figure 2A, B). This was consistent across subjects, as shown by the distributions of correlation coefficients (Figure 2C, D) and the percentage of patient’s that displayed significant positive correlations between aperiodic exponent and CF (Figure 2E).

**Figure 2.**
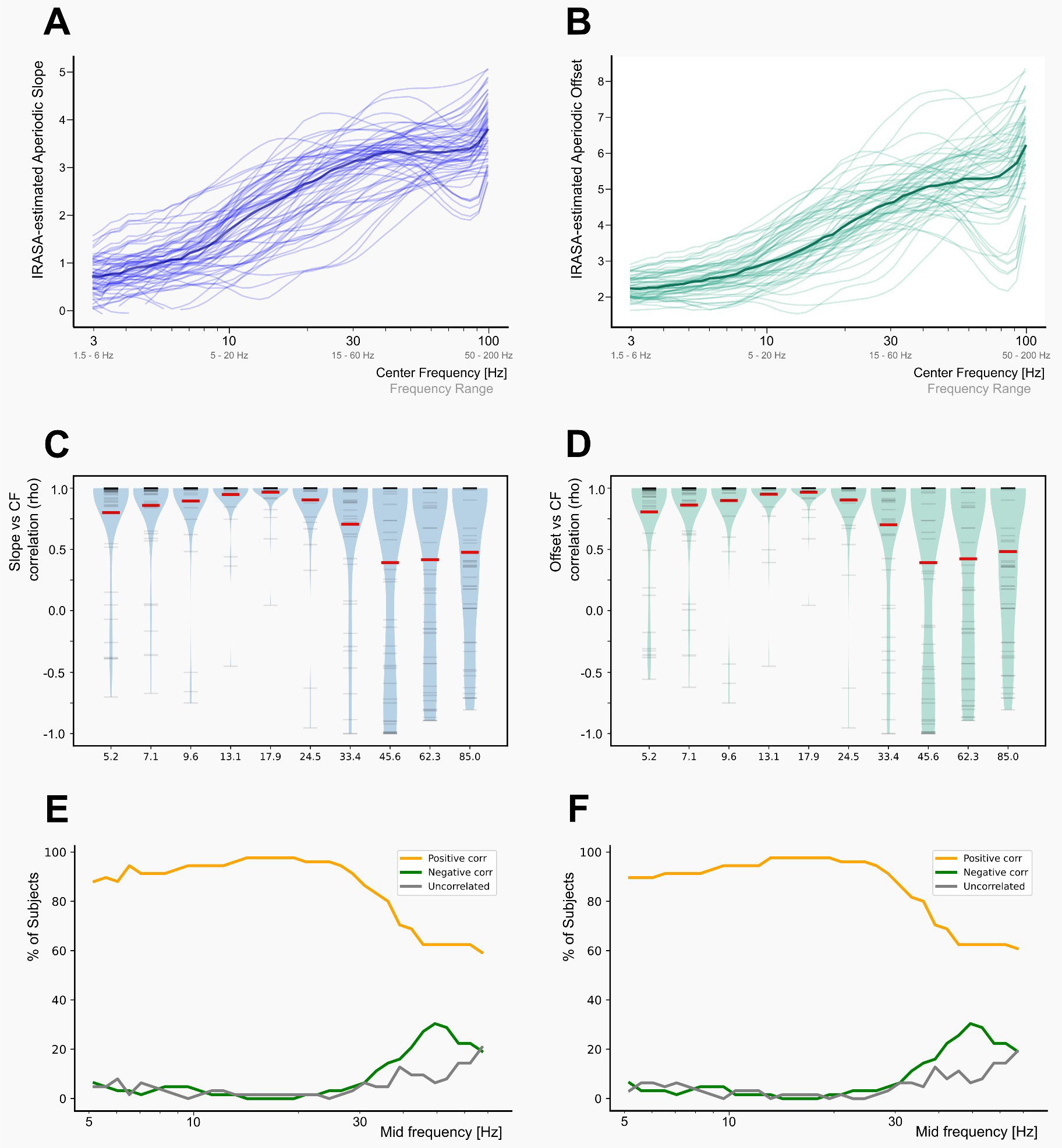
Aperiodic parameters depend on the frequency range in which they are estimated when using IRASA. **(A-B)** Line plot showing, for each subject (soft lines), the dependency of the aperiodic slope (A) and offset (B) versus frequency range (CF) used to estimate them. (**C-D**) Plots illustrating the Spearman correlation coefficients (rho) between 15 aperiodic slopes (C) and the CF of the frequency ranges they were evaluated. Mid frequency refers to the middle frequency of the 15 consecutive frequency ranges. Each subject’s rho values, for each mid frequency is represented by a horizontal grey line in each column. Each column’s violin plot displays the probability density function of rho values across subjects. Median rho values for each mid frequency are represented by a thick red horizontal line in each column. (**E-F**) Percentage of patients that showed either positive, negative or non-significant correlations between aperiodic exponent (E) and CF and between aperiodic exponent (F) and CF for each mid frequency employed. Significance of Spearman correlations was corrected for multiple comparisons.

### iDFT model signals of two unique aperiodic slopes

We observed a continuous dependency between the estimated aperiodic slopes and CF. However, this could be the result of the smoothing of the aperiodic estimates across CFs due to the fact that estimations need to be done in a (relatively wide) frequency range. Thus, a “moving average” effect could be taking place. To evaluate this possibility we made aperiodic estimations in 6 sets of simulated signals, each one built with two power-laws (aperiodic exponents), one in low and one in high frequency ranges. These two power laws changed at 10 Hz (Figure 3 A,B). All sets of simulations had aperiodic slopes of 1 below 10 Hz and varying aperiodic slopes for frequencies above 10 Hz (Figure 3A, B). We estimated aperiodic slopes using Specparam and IRASA methods in these simulated signals and found that indeed smoothing of the aperiodic slope vs CF function was present for both estimation strategies, (Figure 3C, D). IRASA showed a higher degree of smoothing than Specparam. However, both sets of curves displayed clear sigmoid shapes, with plateaus in low and high frequencies instead of a continuous dependency between slope and CF, as seen with empirical data (Figures 1, 2).

**Figure 3.**
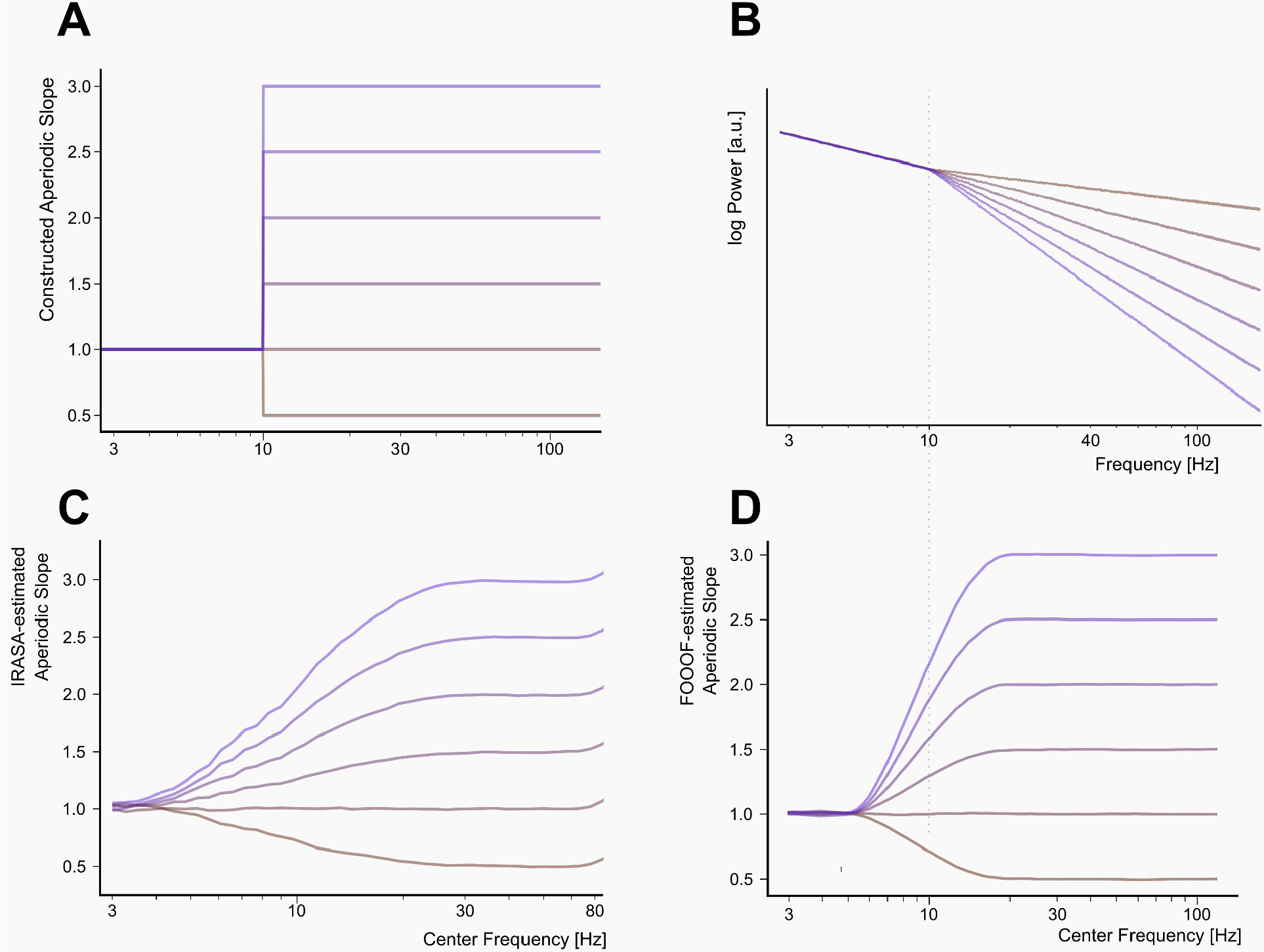
Estimating aperiodic slope with Specparam and IRASA in modeled signals. **(A)** Line plot showing the 6 slopes configurations employed to simulate signals using inverse Discrete Fourier Transform (iDFT) models. All signals have an aperiodic slope of 1 for frequencies lower than 10 Hz and various different aperiodic slopes for frequencies greater than 10 Hz ranging from 0.5 to 3 in increments of 0.5. **(B)** Average PSDs of modeled signals constructed with different aperiodic slopes configurations. **(C)** IRASA estimations of aperiodic slopes calculated across different CFs. **(D)** Specparam estimations of aperiodic slope for each model. The color of each trace is consistent for all panels. Parameters for aperiodic slope estimations were the same as those employed for analyses of empirical data.

## Discussion

In this work we employed high-resolution intracortical human data to systematically evaluate the relation between the aperiodic exponent and the frequency at which it is estimated. Two distinct methods convergently showed that the magnitude of the aperiodic exponent is strongly dependent on frequency range. Specifically, higher frequencies tend to yield greater aperiodic exponents. This dependency was highly consistent across subjects for frequency ranges centered below ∼30 Hz (e.g. 1.5 – 6 Hz to 15 – 60 Hz). We also show that aperiodic offset closely follows the aperiodic exponent across frequencies in our resting-state data. Finally, our results illustrate that the dependency of aperiodic exponent towards frequency can be roughly solved using two power-laws, one for low frequencies and one for high frequencies. However, in order to precisely describe the underlying phenomena of non-oscillatory activity in brain field potentials, a multifractal approach will likely be required.

Power law behaviour has been described for a variety of distinct phenomena from various disciplines, from linguistics (Zipf, 1935) to gravitational waves (Gogoi & Dev Goswami, 2022) and many others (Clauset et al., 2009). They usually fit the data only in a limited range of magnitudes (Stumpf & Porter, 2012). Full power laws can be synthetically generated using few assumptions in surprisingly simple models (Kramer & Chu, 2023; Thurner et al., 1997). In the empirical case of brain field potentials (e.g. EEG), the aperiodic exponent of its power spectral density has been linked to the balance between excitatory and inhibitory neural activity (Duma et al., 2024; Gao et al., 2017; Wiest et al., 2023), a key property of cortical systems. However, a recent report using precise pharmacological interventions, has put into question the universality of the relation between aperiodic slope and E/I balance (Salvatore et al., 2024). Our results are agnostic about this debate, however, they raise a related question. Namely if aperiodic exponents calculated in different frequency ranges (e.g. low and high) are related in the same way to E/I balance, or if they are differentially associated to this and/or to other system-level properties of cortical systems. In other words, it remains an open question whether the aperiodic exponent in low frequencies is a reflection of the same cortical properties as the aperiodic exponents derived from high frequencies.

In light of the general relation between aperiodic exponent and frequency observed here, we wondered if there is a range of frequencies in which the aperiodic exponent is stable across frequencies. We found that there is a continuous, significant and positive correlation between aperiodic exponent and frequency range (Figure 1C, 2C) in most studied subjects (Figure 1E, 2E) when CF was lower than ∼30 Hz. This relatively low frequency range is of particular importance considering that EEG studies that evaluate aperiodic exponents normally only employ frequencies below 50 Hz (Kramer & Chu, 2023). More inconsistent results were found at higher frequencies, which could be in part related to artifacts. We believe, however, that it is important to emphasize the robustness of our results considering the relatively high number of patients and the fact that by the nature of intracortical recordings several low frequency (e.g. sudoration) and high frequency artifacts (e.g. muscular activity) are markedly reduced or absence in comparison to, for example, EEG.

In recent years there has been an increase in aperiodic activity-related research, in particular evaluating EEG’s aperiodic exponent as a possible marker to be used in clinical applications (Donoghue, 2024; Pani et al., 2022). In this context, our results illustrate that aperiodic exponents should not be compared when obtained with different methods (e.g. IRASA or Specparam) or with different frequency ranges (e.g. low vs high), as it could cause erroneous conclusions. To avoid this, it could be argued that a generally agreed upon frequency range to estimate aperiodic activity should be established. In this regard we would tentatively argue that, for EEG signals, a lower limit at or above 12 Hz would avoid the frequently prominent activity in the alpha range and an upper limit below 50 Hz would avoid known artifact sources. However, we believe much more clinical and basic research is required, exploring aperiodic activity in various frequency ranges, before a solid conclusion could be reached in this matter.

## References

Buzsáki, G., Anastassiou, C. A., & Koch, C. (2012). The origin of extracellular fields and currents— EEG, ECoG, LFP and spikes. Nature Reviews Neuroscience, 13(6), 407–420. 10.1038/nrn3241

Clauset, A., Shalizi, C. R., & Newman, M. E. J. (2009). Power-Law Distributions in Empirical Data. SIAM Review, 51(4), 661–703. 10.1137/070710111

Coa, R., La Cava, S. M., Baldazzi, G., Polizzi, L., Pinna, G., Conti, C., Defazio, G., Pani, D., & Puligheddu, M. (2022). Estimated EEG functional connectivity and aperiodic component induced by vagal nerve stimulation in patients with drug-resistant epilepsy. Frontiers in Neurology, 13, 1030118. 10.3389/fneur.2022.1030118

Donoghue, T. (2024). A systematic review of aperiodic neural activity in clinical investigations. Psychiatry and Clinical Psychology. 10.1101/2024.10.14.24314925

Donoghue, T., Haller, M., Peterson, E. J., Varma, P., Sebastian, P., Gao, R., Noto, T., Lara, A. H., Wallis, J. D., Knight, R. T., Shestyuk, A., & Voytek, B. (2020). Parameterizing neural power spectra into periodic and aperiodic components. Nature Neuroscience, 23(12), 1655–1665. 10.1038/s41593-020-00744-x

Donoghue, T., Hammonds, R., Lybrand, E., Washcke, L., Gao, R., & Voytek, B. (2024). Evaluating and Comparing Measures of Aperiodic Neural Activity. Neuroscience. 10.1101/2024.09.15.613114

Duma, G. M., Cuozzo, S., Wilson, L., Danieli, A., Bonanni, P., & Pellegrino, G. (2024). Excitation/Inhibition balance relates to cognitive function and gene expression in temporal lobe epilepsy: A high density EEG assessment with aperiodic exponent. Brain Communications, 6(4), fcae231. 10.1093/braincomms/fcae231

Ferree, T. C., & Hwa, R. C. (2003). Power-law scaling in human EEG: Relation to Fourier power spectrum. Neurocomputing, 52–54, 755–761. 10.1016/S0925-2312(02)00760-9

Gao, R., Peterson, E. J., & Voytek, B. (2017). Inferring synaptic excitation/inhibition balance from field potentials. NeuroImage, 158, 70–78. 10.1016/j.neuroimage.2017.06.078

Gerster, M., Waterstraat, G., Litvak, V., Lehnertz, K., Schnitzler, A., Florin, E., Curio, G., & Nikulin, V. (2022). Separating Neural Oscillations from Aperiodic 1/f Activity: Challenges and Recommendations. Neuroinformatics, 20(4), 991–1012. 10.1007/s12021-022-09581-8

Gogoi, D. J., & Dev Goswami, U. (2022). Gravitational waves in f(R) gravity power law model. Indian Journal of Physics, 96(2), 637–646. 10.1007/s12648-020-01998-8

He, B. J., Zempel, J. M., Snyder, A. Z., & Raichle, M. E. (2010). The Temporal Structures and Functional Significance of Scale-free Brain Activity. Neuron, 66(3), 353–369. 10.1016/j.neuron.2010.04.020

Kramer, M. A., & Chu, C. J. (2023). The 1/f-like behavior of neural field spectra are a natural consequence of noise driven brain dynamics. 10.1101/2023.03.10.532077

Kucyi, A., Daitch, A., Raccah, O., Zhao, B., Zhang, C., Esterman, M., Zeineh, M., Halpern, C. H., Zhang, K., Zhang, J., & Parvizi, J. (2020). Electrophysiological dynamics of antagonistic brain networks reflect attentional fluctuations. Nature Communications, 11(1), 325. 10.1038/s41467-019-14166-2

Lanzone, J., Colombo, M. A., Sarasso, S., Zappasodi, F., Rosanova, M., Massimini, M., Di Lazzaro, V., & Assenza, G. (2022). EEG spectral exponent as a synthetic index for the longitudinal assessment of stroke recovery. Clinical Neurophysiology, 137, 92–101. 10.1016/j.clinph.2022.02.022

Lendner, J. D., Helfrich, R. F., Mander, B. A., Romundstad, L., Lin, J. J., Walker, M. P., Larsson, P. G., & Knight, R. T. (2020). An electrophysiological marker of arousal level in humans. eLife, 9, e55092. 10.7554/eLife.55092

Medel, V., Irani, M., Crossley, N., Ossandón, T., & Boncompte, G. (2023). Complexity and 1/f slope jointly reflect brain states. Scientific Reports, 13(1), 21700. 10.1038/s41598-023-47316-0

Mostile, G., Giuliano, L., Dibilio, V., Luca, A., Cicero, C. E., Sofia, V., Nicoletti, A., & Zappia, M. (2019). Complexity of electrocortical activity as potential biomarker in untreated Parkinson’s disease. Journal of Neural Transmission, 126(2), 167–172. 10.1007/s00702-018-1961-6

Newman, M. (2005). Power laws, Pareto distributions and Zipf’s law. Contemporary Physics, 46(5), 323–351. 10.1080/00107510500052444

Ostlund, B. D., Alperin, B. R., Drew, T., & Karalunas, S. L. (2021). Behavioral and cognitive correlates of the aperiodic (1/f-like) exponent of the EEG power spectrum in adolescents with and without ADHD. Developmental Cognitive Neuroscience, 48, 100931. 10.1016/j.dcn.2021.100931

Pani, S. M., Saba, L., & Fraschini, M. (2022). Clinical applications of EEG power spectra aperiodic component analysis: A mini-review. Clinical Neurophysiology, 143, 1–13. 10.1016/j.clinph.2022.08.010

Pereda, E., Gamundi, A., Rial, R., & González, J. (1998). Non-linear behaviour of human EEG: Fractal exponent versus correlation dimension in awake and sleep stages. Neuroscience Letters, 250(2), 91–94. 10.1016/S0304-3940(98)00435-2

Robertson, M. M., Furlong, S., Voytek, B., Donoghue, T., Boettiger, C. A., & Sheridan, M. A. (2019). EEG power spectral slope differs by ADHD status and stimulant medication exposure in early childhood. Journal of Neurophysiology, 122(6), 2427–2437. 10.1152/jn.00388.2019

Salvatore, S. V., Lambert, P. M., Benz, A., Rensing, N. R., Wong, M., Zorumski, C. F., & Mennerick, S. (2024). Periodic and aperiodic changes to cortical EEG in response to pharmacological manipulation. Journal of Neurophysiology, 131(3), 529–540. 10.1152/jn.00445.2023

Steriade, M. (2005). Cellular Substrates of Brain Rhythms. In Electroencephalography: Basic Principles, Clinical Applications, and Related Fields (Fifth Edition). Lippincott Williams and Wilkins.

Stumpf, M. P. H., & Porter, M. A. (2012). Critical Truths About Power Laws. Science, 335(6069), 665–666. 10.1126/science.1216142

Thurner, S., Feurstein, M. C., & Teich, M. C. (1997). Conservation laws in coupled multiplicative random arrays lead to $1/f$ noise (No. arXiv:adap-org/9709005). arXiv. http://arxiv.org/abs/adap-org/9709005

Wen, H., & Liu, Z. (2016). Separating Fractal and Oscillatory Components in the Power Spectrum of Neurophysiological Signal. Brain Topography, 29(1), 13–26. 10.1007/s10548-015-0448-0

Wiest, C., Torrecillos, F., Pogosyan, A., Bange, M., Muthuraman, M., Groppa, S., Hulse, N., Hasegawa, H., Ashkan, K., Baig, F., Morgante, F., Pereira, E. A., Mallet, N., Magill, P. J., Brown, P., Sharott, A., & Tan, H. (2023). The aperiodic exponent of subthalamic field potentials reflects excitation/inhibition balance in Parkinsonism. eLife, 12, e82467. 10.7554/eLife.82467

Zhang, Y., Wang, Y., Cheng, H., Yan, F., Li, D., Song, D., Wang, Q., & Huang, L. (2023). EEG spectral slope: A reliable indicator for continuous evaluation of consciousness levels during propofol anesthesia. NeuroImage, 283, 120426. 10.1016/j.neuroimage.2023.120426

Zipf, G. K. (1935). The Psycho-biology of Language An Introduction to Dynamic Philology (The University of Michigan). Mifflin.

